# Disruption of spindle orientation and protein localization during asymmetric cleavage by pharmacological inhibition of serotonin signaling

**DOI:** 10.1101/2025.04.19.649612

**Authors:** Ayaki Nakamoto, Lisa M. Nagy

## Abstract

Cell polarity directs the orientation and size of asymmetric cell divisions, and the segregation of cell fate determinants processes fundamental to development in all multicellular organisms. During asymmetric cleavage, the mitotic spindle aligns with a specified polarity of the mother cell, and cell fate determinants are distributed asymmetrically along the division axis. Here, we report that pharmacological inhibition of serotonin signaling during the 4-to-8-cell division in early embryos of the mud snail *Ilyanassa obsoleta* (currently known as *Tritia obsoleta*) disrupts the typical unequal division pattern. The oblique axis of division common to spirally cleaving molluscan embryos is altered, and the position of the mitotic spindle is randomized in these treatments. Mother cells generate abnormally large, atypically positioned daughter cells. We also find that, in normal embryos, proteins recognized by phosphorylated PKC and Bazooka/PAR-3 antibodies typically co-localize with the spindle apparatus to the apical cortex of each mother cell. These antigens subsequently segregate to the smaller of the two daughter cells. In embryos treated with the serotonin-receptor antagonist, the localization of these asymmetrically segregating proteins is randomized, and their localization is independent of spindle position. These results suggest that serotonin signaling coordinates spindle orientation, cortical polarity, and cell size in early asymmetric cleavages.

**Highlights:**

- This study suggests a novel rule for serotonin signaling in asymmetric cleavage
- Serotonin-receptor antagonist treatment disrupts typical asymmetric cleavage
- aPKC and Bazooka/PAR-3 localize with the spindle at the apical cortex
- Inhibition of serotonin signaling disrupts localization of the polarity proteins
- Serotonin may regulate division through G-protein effects on the cytoskeleton

## 1. Introduction

Asymmetric cleavage is essential for generating cellular diversity during development. Generally, the mother cell is polarized along a specific axis by external cues or localized intrinsic factors, and the mitotic spindle is oriented according to the polarized signal. Specific mRNAs and proteins are segregated into one daughter cell, which then follows distinct cell fates. In the embryos of model organisms such as vertebrates, flies, and nematodes, spindle orientation is regulated by an evolutionally conserved set of polarity proteins (e.g., atypical PKC, PAR-6, Bazooka/PAR-3). The proteins localize to specific cortical domains, where they associate with other molecular players (e.g., Pins/LGN, NuMA) to regulate spindle orientation (Gönczy, 2008; Suzuki, 2006; Lu and Johnston, 2013; di Pietro et al., 2016).

In a number of lophotrochozan species, the early embryos undergo asymmetric cleavages known as spiral cleavage (Conklin, 1897; Crampton, 1896; Freeman and Lundelius, 1992; Lambert, 2010; Henry, 2014; Martín-Durán and Marlétaz; 2020). During spiral cleavage, each cell (macromere) at the four-cell stage functions as a stem cell and undergoes a series of asymmetric divisions, each of which generates a significantly smaller daughter cell (micromere) (Fig. 1A). During these divisions, the spindle axis is directionally polarized at an oblique angle to the animal-vegetal axis of the embryo. In subsequent division cycles, the division angle alternates between clockwise and counterclockwise with respect to the animal pole. These stem-cell divisions also lead to asymmetric inheritance of cell-fate determinants, which depends on an intact actin filament network (Lambert and Nagy, 2002; Kingsley et al. 2007). Whether the spindle orientation in spiral cleavage depends on the polarity proteins conserved in model systems has not been previously reported.

**Figure 1.**
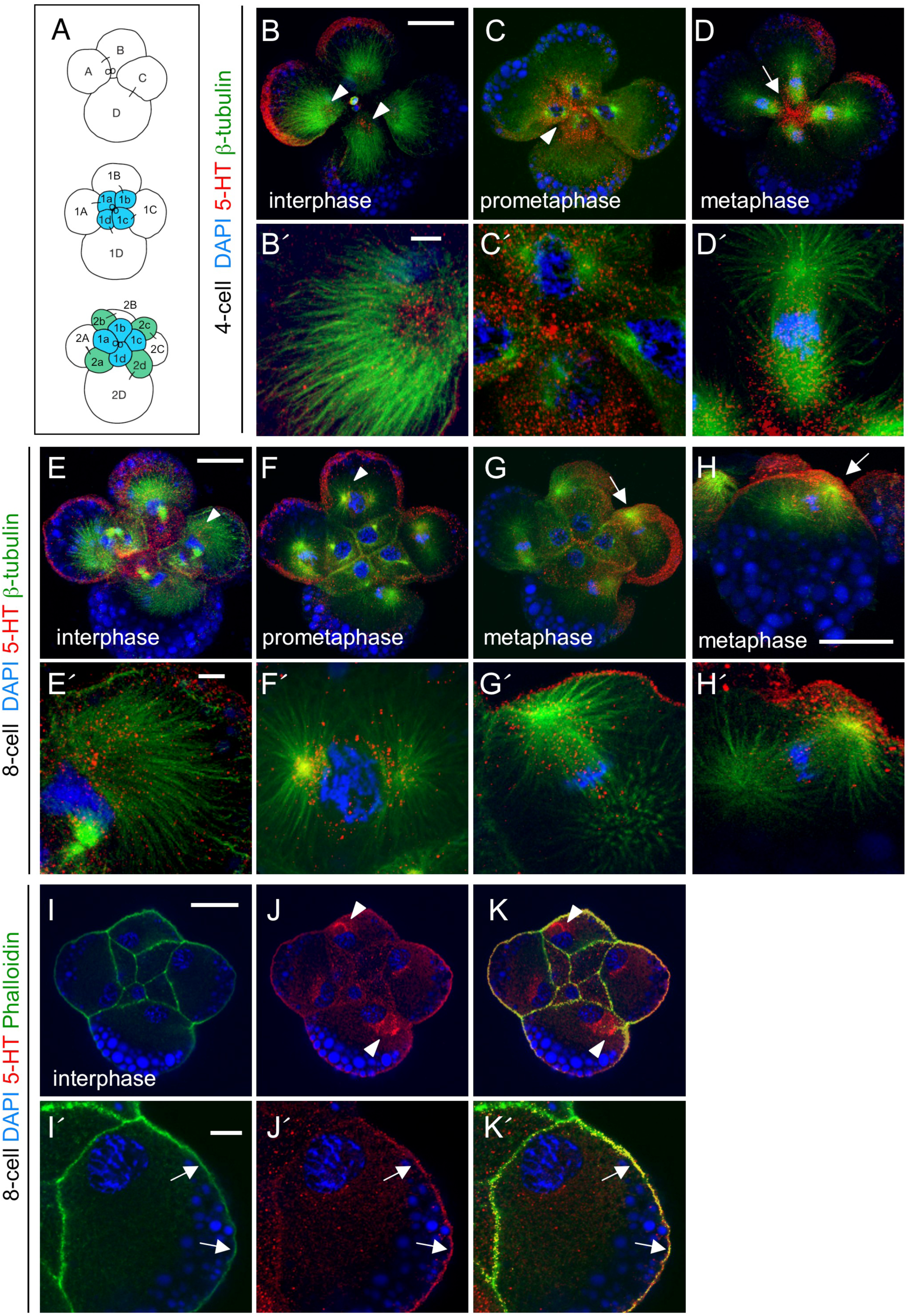
Early asymmetric cleavage and expression of serotonin in early *Ilyanassa* embryos. (A) Schematic figures of the first three asymmetric macromere divisions (4, 8 and 12-cell stages) of the *Ilyanassa* embryo [adapted from Lambert and Nagy (2002)]. Four macromeres (A-D) are generated during the first two division cycles. During each successive cleavage cycle, each macromere (stem cell) undergoes an asymmetric division and produces a smaller daughter cell (micromere) with a distinct cell fate. The macromeres (A-D) divide at an oblique angle (less than 90°), clockwise from the animal-vegetal axis, and generate the first daughter cells (blue). In the next cleavage, macromeres (1A-1D) divide at a similar oblique angle, counterclockwise from the animal-vegetal axis, and generate the second daughter cells (green). (B-H′) Serotonin expression during the 4 (B-D′) and 8-cell stages (E-H′). Co-staining with serotonin (red) and β-tubulin (green) antibodies and DAPI stained nuclei (blue) are shown. (B-G′) Animal pole views. (H, H′) Side views of 1D macromere. Animal pole is to the top. During the interphase of the 4 (B, B′) and 8-cell stages (E, E′), serotonin was detected in the center of the array of microtubules (arrowheads) and the macromere cortex. During prometaphase (C, C′; F, F′), an apical section shows that serotonin is concentrated in the apical cytoplasm (C, arrowhead). In a deeper section, serotonin is observed in the perinuclear region of the macromere (F, arrowhead). During metaphase, serotonin is localized to the surface of macromere (D, G, H arrow) and concentrated around the apical spindle pole (D′, G′, H′). (I-K′) Co-staining with phalloidin (green) and serotonin antibody (red). Nuclei (blue) were stained with DAPI. Serotonin is localized in a spherical structure next to the nuclei (J, arrowheads) and cortex of the macromere. Note that phalloidin and serotonin signals overlap in the cortex (arrows). Scale bars, 50 µm (A-K); 10 µm (A′-K′).

Some insight into the regulation of spindle orientation during spiralian development has emerged from studies in the freshwater snail *Lymnaea stagnalis*, where the chirality of shell coiling depends on the position of the mitotic spindle during the third cleavage. Treatments with depolymerizing drugs such as latrunculin showed that the asymmetric division of the macromeres depends on an intact actin cytoskeleton (Shibazaki et al., 2004). More recently, it was discovered that the diaphanous-related gene (referred to as *Lsdia2* by Davison et al., 2016 and *Lsdia1* by Kuroda et al., 2016) determines whether the division pattern is clockwise (resulting in a dextral-coiling shell) or counterclockwise (resulting in a sinistral-coiling shell; Davison et al., 2016; Kuroda et al., 2016; Abe and Kuroda 2019). However, the embryos of the homozygous knockout of *Lsdia1* underwent a typical spiral cleavage pattern (Abe and Kuroda, 2019). Thus, the spiral cleavage pattern is generated independently of *Lsdia1,* leaving the molecular mechanisms that initially polarize the spindle in spiral cleavage yet to be discovered.

In many circumstances, the polarization of the mitotic spindle depends on cell-cell signaling. Common examples include Wnt and Notch signaling (e.g. Henry et al., 2008; Habib et al., 2013; Bhat, 2014). The pre-neural activity of serotonin (5-hydroxytryptamin, 5-HT) signaling has also been implicated in regulating early embryonic cell divisions. Serotonin antagonist treatment disrupts unequal cleavage in sea urchin embryos (Shmukler, 2010) and inhibits the first cleavage divisions in the mollusk *Tritonia* and annelid *Ophryotrocha*, (Buznikov et al., 2003; Emanuelsson, 1974; Emanuelsson, 1992). Overexpression of the serotonin precursor 5-HTP disrupts cell arrangements in the *Lymnaea* blastula (Bogomolov and Voronezhskaya, 2022). A growing body of literature suggests that serotonin is explicitly used during embryogenesis to regulate left-right asymmetry, cell proliferation, differentiation, and cell movements (Buznikov et al., 2001; Buznikov and Shmukler, 1981; Fukumoto et al., 2005; Shmukler Iu, 2010; Karki et al., 2023). Whether the observed effects of loss or overexpression of serotonin in these cases are a consequence of disruption of the mitotic spindle has not been explored.

In this study, we examined whether serotonin signaling is involved in proper spindle orientation and protein localization during early asymmetric cleavages in the mud snail *Ilyanassa obsoleta* (currently *Tritia obsoleta*). We characterized the expression of serotonin and examined the role of serotonin signaling in early *Ilyanassa* embryos. When a serotonin-receptor antagonist inhibited serotonin signaling, macromeres divided along abnormal orientations and generated significantly large micromeres. We also found that antibodies for the phosphorylated PKC isoforms and Bazooka/PAR-3 (Wodarz et al., 1999) recognize antigens that localize to the specific apical cortex of macromeres (i.e., prospective spindle-pole attachment site), indicating that this molecular machinery may be conserved in *Ilyanassa* embryos. When serotonin signaling was inhibited, these proteins localized to ectopic positions, and the relationship between the protein localization and spindle orientation was disrupted. Taken together, our results suggest that serotonin signaling is required for coordinating proper spindle orientation, protein localization, and cell size, providing a novel insight into the regulation of asymmetric cleavage.

## 2. Results and Discussion

### 2.1. Expression of serotonin and serotonin receptors during the embryonic stages

In the early *Ilyanassa* embryo, serotonin expression exhibited dynamic, cell-cycle-dependent expression throughout the first four cleavage stages, showing co-localization with cytoskeletal elements in the developing spindle poles and cell cortex (Fig. 1B-K′ and Supplementary Fig. S1). Cytoplasmic and membrane-associated serotonin were detected at the one- and two-cell stages (Supplementary Fig. S1), consistent with maternal provision of the peptide. During interphase, serotonin was detected in the microtubule array’s center and throughout each cell’s cortex (Fig. 1B, B′; 1E, E′). In prometaphase, serotonin became concentrated in the apical cytoplasm (Fig. 1C, C′) and was detected in the perinuclear region associated with the developing spindle pole (Fig. 1F, F′). Serotonin subsequently became enriched at the apical spindle pole during metaphase (Fig. 1D, D′; G, G′; H, H′). This expression pattern suggests a dynamic correlation between serotonin and the mitotic spindle pole. In addition, serotonin co-localized with phalloidin at the cortex, consistent with a close association between serotonin and the actin cytoskeleton (Fig. 1I-K′). To control for the specificity of the antibody, the anti-5-HT antibody was pre-incubated with 5-HT-BSA conjugate (ImmunoStar). No specific staining was observed after the depletion of the antibody (Supplementary Fig. S2). We also identified orthologs of the serotonin receptors (5-HT1 and 5-HT2) in an *Ilyanassa* EST database (Lambert et al., 2010) (Supplemental Fig. S3). Both 5-HT1 and 5-HT2 mRNAs were detected in 4-12 cell embryos using RT-PCR (Supplemental Fig. S4A). Expression of the serotonin receptor protein in early embryos was also confirmed via Western blot and mass spectrometry (Supplemental Fig. S4B-D). Based on these results, we conclude that both serotonin and serotonin receptor proteins are expressed in the early *Ilyanassa* embryo.

### 2.2 Disruption of asymmetric cleavage by serotonin receptor antagonist

To analyze the function of serotonin signaling, we applied the highly selective antagonist metergoline (Sigma), commonly used to block serotonin signaling in various model organisms including mammals (Beretta et al., 1965), nematodes (Hamdan et al., 1999), and molluscs (Cohen et al., 2003; Sugamori et al., 1993). In this study, we focused on the division of the macromeres at the 8-cell stage. In this division, the macromere divides asymmetrically (Fig. 2A, A′): the size ratio of the daughter/mother cell (micromere/macromere) is 0.26 ± 0.09 (Fig. 2G; *n*=37). When embryos were treated with 30 µM metergoline from the early 4-cell stage to the early 8-cell stage, macromeres divided nearly equally (Fig. 2C, C′), or they underwent somewhat unequal division, in which the larger size of the daughter cell was readily apparent due to abnormal inheritance of yolk (arrow in Fig. 2C). The micromere/macromere size ratio was significantly greater than untreated embryos (Fig. 2G; 0.81 ± 0.19, *n*=45, Student’s t-test; p<0.0001), suggesting that the division plane had shifted to a more symmetrical position.

**Figure 2.**
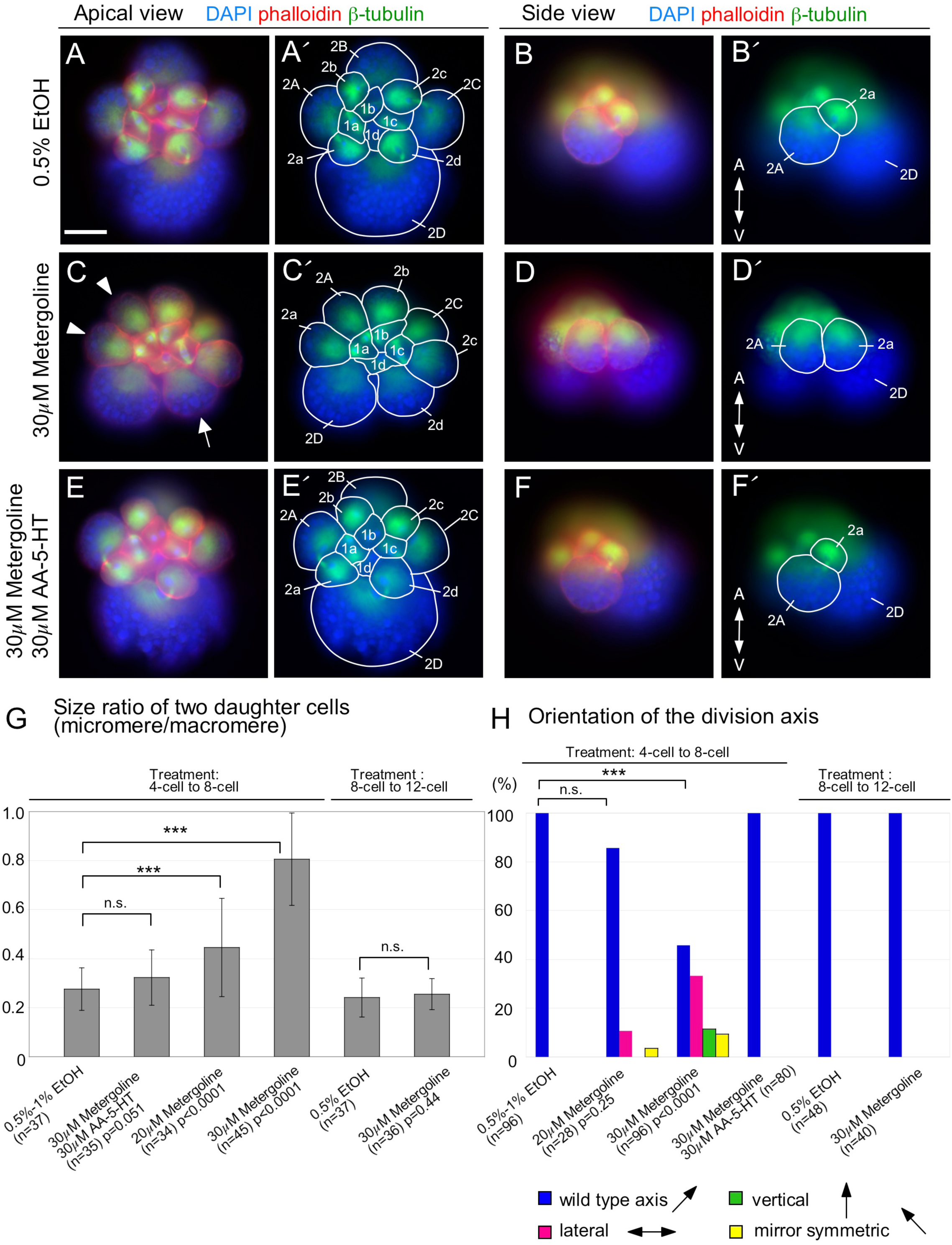
Serotonin receptor antagonist disrupts asymmetric cleavage. (A-F′) Phenotypes of control and drug-treated embryos. β-tubulin antibody expression (green), phalloidin (red), and DAPI-stained nuclei (blue) are shown. (A, C, E) apical view. (B, D, F) side view. A; animal pole, V; vegetal pole. In the corresponding images (A′-F′), the cell boundaries are outlined, and the cells are identified. (A-B′) Control (0.5%EtOH) treatment. Macromeres produce micromere progeny along an oblique orientation (B, B′) with respect to the animal-vegetal axis. (C-D′) 30 µM metergoline treatment. Macromeres divide equally (C, arrowheads) or undergo somewhat unequal cleavage in which the yolk abnormally segregates (C, arrow). In addition, macromeres divide laterally with respect to the animal-vegetal axis (D, D′). The daughter cells produced from an equal macromere cleavage were designated based on their relative position to the D macromere. In C and C′, the 2B cell is out of focus because of the vertical division of the mother cell. (E, E′) Co-treatment of metergoline and AA-5-HT. Macromeres divide normally and embryos are indistinguishable from controls (A-B′). (G) The size ratio of two daughter cells (micromere/macromere). Statistical significance was analyzed using the Student’s t-test; n. s.; not significant; ***, p<0.0001; data are presented as means ± s.e.m. (H) Orientation of the division axis. Statistical significance was analyzed by the Mann-Whitney test; n. s.; not significant; ***, p<0.0001.

We also found that the division axis of the macromere failed to orient correctly in metergoline-treated embryos (Fig. 2; 54%, *n*=52/96, Mann-Whitney test; p<0.0001). In controls (0.5%-1% EtOH), the macromeres always divided along a specific oblique orientation with respect to the animal-vegetal axis (*n*=96, Fig. 2B, B′). However, when treated with 30 µM metergoline, 33% (*n*=32/96) divided laterally (Fig. 2D, D′) and 12% (*n*=11/96) divided vertically. 20 µM metergoline treatment had milder effects on the size ratio of the daughter cells (0.45 ± 0.20, *n*=34, Student’s t-test; p<0.0001), although the effect of 20 µM metergoline on the division axis was not significant (14%, *n*=4/28, Mann-Whitney test; p<0.5; 11% (*n*=3/28) divided laterally, 4% (*n*=1/28) divided vertically). These results suggest that the effect of metergoline is dose dependent. When embryos were co-treated with 30 µM metergoline and 30 µM arachidonyl serotonin (AA-5-HT, Tocris), a lipophilic and membrane-permeable agonist of 5-HT (Buznikov et al., 2001; Buznikov et al., 2003) during the 4-8 division, all the mitotic spindles in the macromeres were oriented typically (Fig. 2 E-F′; *n*=80) and the size ratio of the daughter cells was similar to those of control treatment (0.32 ± 0.11, *n*=35, Student’s t-test; p=0.051). The resultant treated embryos were indistinguishable from control embryos. Thus, the phenotype of metergoline treatment could be rescued by co-treatment with the agonist AA-5-HT.

Embryos are sensitive to the antagonist only during the first asymmetric cleavage they undergo, from the early 4-cell to the early 8-cell stage. When embryos were treated with 30 µM metergoline from the early 8-cell stage to the early 12-cell stage, all the macromeres divided normally (*n*=40 for the division axis and *n*=36 for the size ratio, Fig. 2G, H). This suggests that serotonin signaling occurs during the third cleavage cycle (4-8 cells, production of first set of micromeres) and this signaling is required for proper fourth cleavage (8-12 cells, production of second set of micromeres). In *Lymnaea stagnalis*, an elegant micromanipulation study has shown that blastomere arrangement at the 8-cell stage determines the orientation of the fourth cleavage and the chirality of the adult shell (Kuroda et al., 2009). Thus, the polarity of the fourth cleavage is determined during (or shortly after) the third cleavage in both *Ilyanassa* and *Lymnaea*.

### 2.3. Expression of polarity marker proteins and their ectopic localization by serotonin receptor antagonist

In many metazoan animals, spindle orientation is regulated by a conserved set of proteins, including atypical PKC and the Par polarity proteins (Gönczy, 2008; Suzuki, 2006; Lu and Johnston, 2013; di Pietro et al., 2016). To determine the relationship between serotonin signaling and spindle orientation, we analyzed the expression of antibodies known to mark spindle polarity in both control and metergoline-treated embryos. We found that the antibody that recognizes the phosphorylated PKC isoforms (pPKC; Cell Signaling Technology; Fig. 3A-E) and the anti-Bazooka/PAR-3 antibody (Wodarz et al., 1999) (Fig. 3 F-H) recognize antigen(s) that localize to the apical cortex of *Ilyanassa* macromeres and micromeres in a cell-cycle dependent manner. During interphase of the 4-cell stage, pPKC-like antigens localize to the centrosome positioned between the nucleus and the apical membrane of each macromere (Fig. 3A). Between prophase and metaphase (Fig. 3B, C), pPKC-like antigens move to the apical cortex that coincides with the apical-spindle-pole microtubules. During telophase (Fig. 3D), the smaller daughter cell inherits the cortically-localized pPKC-like antigens. (Details of the expression pattern are in Supplemental Fig. S5). Similar cell-cycle-dependent expression was observed during the 8-cell stage (Fig. 3I-L). When embryos were treated with metergoline from the early 4-cell stage to the early 8-cell stage, pPKC-like antigens were detected on the centrosomes; however, the centrosomes remained far from the apical membrane in the macromeres (Fig. 3M, N). During prometaphase and metaphase, pPKC-like antigens failed to migrate to the apical macromere cortex and spindles failed to attach to the apical cortex (Fig. 3O, P). Thus, disruption of serotonin signaling also disrupts the localization of proteins typically associated with the asymmetric localization of the spindle pole.

**Figure 3.**
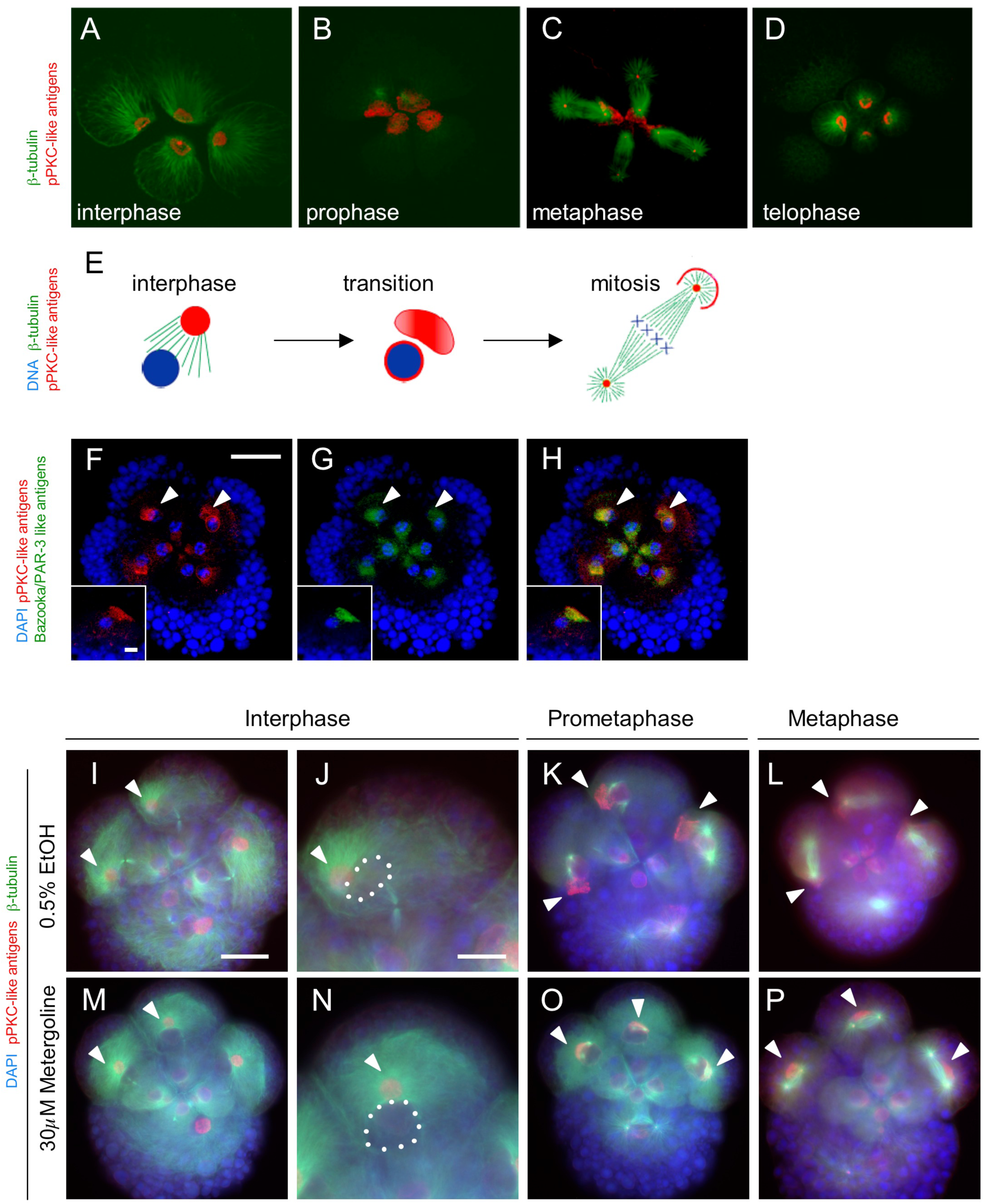
Expression of polarity marker proteins and disruption of their localization by Metergoline treatment. (A-D) Expression of pPKC-like antigens (red) and β-tubulin (green) during the four-to eight-cell division. Apical view. pPKC-like antigens are localized in the interphase centrosome (A). During prophase (B) and prometaphase (C), pPKC-like antigens migrate to the apical cortex of the macromere. pPKC-like antigens are also present in the center of both ends of all four spindles (C). During the telophase (D), pPKC-like antigens are segregated into daughter micromeres. (E) Schematic summary of the pPKC-like antigen expression. See text and Fig. S5 for details. (F-H) Expression of pPKC-like antigens (F) and Bazooka/PAR-3-like antigens (G). Apical view of the 8-cell stage. Bazooka/PAR-3-like antigens (green) are co-localized with pPKC-like antigens (red) of the macromeres (arrowheads). Insets show a lateral view of a macromere of the different embryos. (I-P) Disruption of spindle orientation and pPKC-like antigen localization by Metergoline treatment. In controls, pPKC-like antigens localize to the macromere interphase centrosome (I, arrowheads). In higher magnification view (J), the centrosome is positioned between nuclei (indicated dotted lines) and the apical membrane. During the prometaphase (K) and metaphase (L), pPKC-like antigens localize to the apical cortex of the macromeres, and spindles are attached to the apical cortex (arrowheads). When embryos are treated with Metergoline, centrosomes are localized at abnormal positions (M, arrowheads). They are far away from the apical cortex (N, arrowhead). During prometaphase (O) and metaphase (P), pPKC-like antigens fail to localize to the apical cortex (arrowheads), and spindles are not attached to the apical cortex. Scale bars, 50 µm (F-H, I, K, L, M, O, P); 25 µm (J, N) 10 µm (F-H, inset).

To determine the extent of correspondence between the spindle attachment site and ectopic localization of pPKC-like antigens, spindle orientation and localization of pPKC-like antigens were measured with reference to the animal-vegetal axis of the embryo (Fig. 4A). In controls, spindle orientation and localization of pPKC-like antigens are oblique with respect to the midline reference axis (56 ± 13°, *n*=24 for spindle orientation, 51 ± 12°, *n*=24 for pPKC-like antigens). There was no significant difference between spindle orientation and localization of pPKC-like antigens (Student’s t-test; p=0.24), supporting our observation that pPKC-like antigens localize to the apical spindle attachment site. In contrast, in metergoline-treated embryos, spindle orientation and localization of the pPKC-like antigens were severely disrupted. The angle between the axis of the mitotic spindle and the reference axis was wider than that of controls (75± 22°, *n*=23, Student’s t-test; p<0.001). In particular, 52% (*n*=12/23) of the spindles were aligned laterally with respect to the reference axis (75°-120°). We confirmed the ectopic localization of pPKC-like antigens (120 ± 38°, *n*=23, Student’s t-test; p<0.0001) and observed that spindle orientation and localization of pPKC-like antigens were not correlated (Student’s t-test; p<0.0001). Thus, the disruption of the proteins that mark cortical polarity and the disruption of spindle orientation were independent phenotypes. Similar results were obtained with the Bazooka/PAR-3-like antigens. In controls, the Bazooka/PAR-3-like antigens co-localized with the pPKC-like antigens at the apical cortex (Figs 3F-H, 4D-F). When embryos were treated with metergoline, Bazooka/PAR-3-like antigens localized to ectopic positions within the macromere (Fig. 4G-I). Notably, Bazooka/PAR-3-like antigens remained co-localized with pPKC-like antigens in metergoline treatments. These results suggest that the apical cortex no longer attracts either of these proteins or the spindle in metergoline treated embryos. These results also suggest that serotonin signaling is involved in coordinating cortical polarity and the spindle axis in early *Ilyanassa* embryos. At present, the specificities of these antibodies are not fully determined in *Ilyanassa*; however, based on the cortical expression patterns of these antigens, it is possible that function of the aPKC and Bazooka/PAR-3 system is conserved in lophotrochozoan embryos.

**Figure 4.**
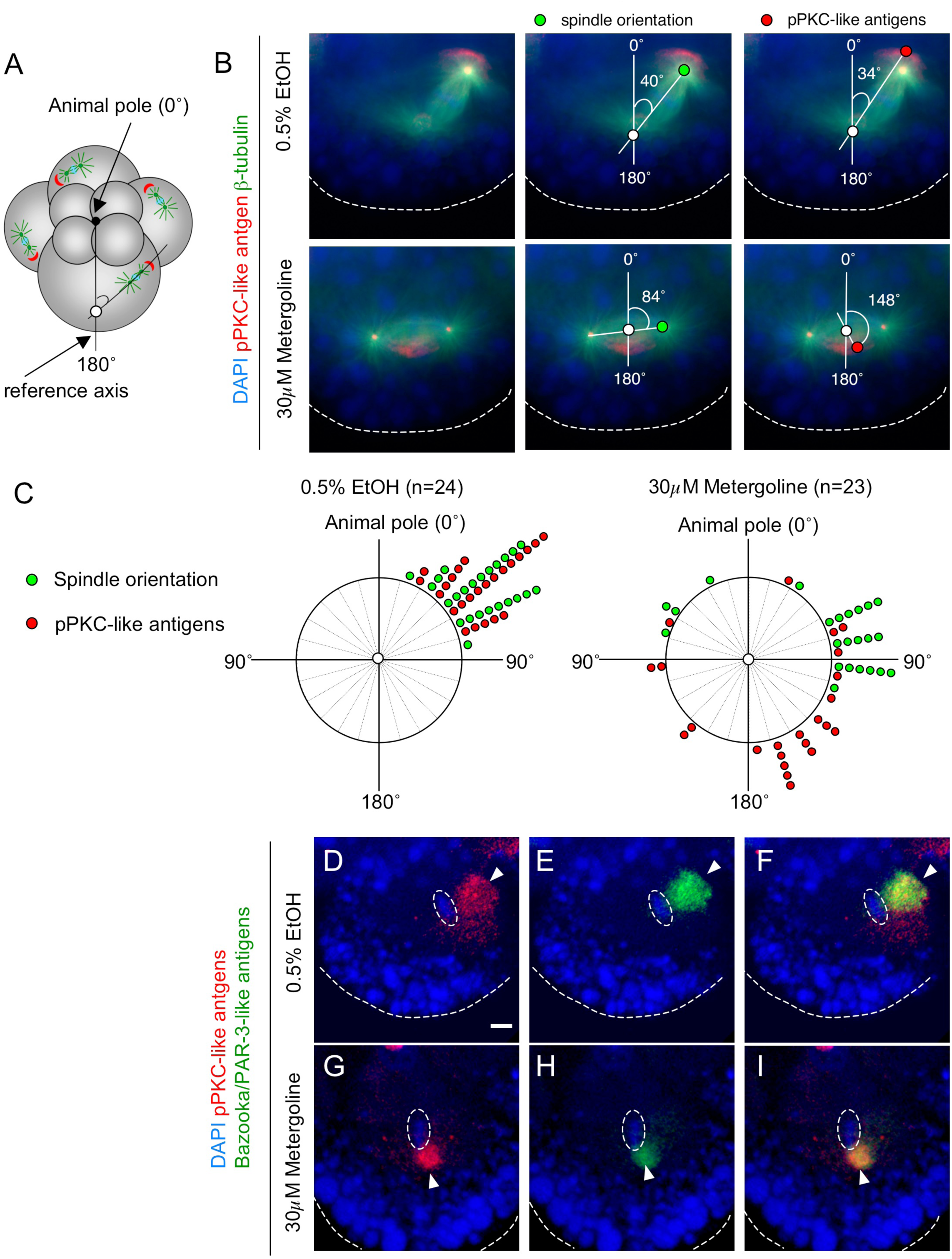
Apical localization of proteins associated with the spindle attachment site is lost in Metergoline-treated embryos. (A) Schematic figure showing the measurements of spindle orientation and pPKC-like antigen localization. The animal pole (0°) is defined as the position where the four micromeres meet. The other point (180°) is defined as the farthest position from the animal pole. The midline axis along these two points is used as a reference axis. (B) Orientation of spindle (green) and localization of pPKC-like antigens (red) in relation to the reference axis. Nuclei (blue) are stained with DAPI. The outline of the macromere is partially indicated with dashed lines. (B, top) Control treatment. (B, bottom) Metergoline treatment. (C) Summary of the angle measurements of spindle orientation (green) and pPKC-like antigens (red). Each circle represents one case. (D-I) Localization of Bazooka/PAR-3 antigen (green) and pPKC-like antigens (red) in the control (D-F) and Metergoline-treated embryo (G-I) at metaphase. Animal pole is to the top. Note that Bazooka/PAR-3 antigen and pPKC-like antigens overlap (arrowheads). Dashed lines circle metaphase chromosomes. Scale bars, 10 µm.

### 2.4. Actin cytoskeleton is required for proper protein localization and division axis

The expression pPKC-like antigens were similar to those of centrosomally localized mRNA during the early asymmetric cleavages (Lambert and Nagy, 2002). It has been shown that movement of the mRNA from the centrosome to the apical cortex requires an intact actin cytoskeleton (Lambert and Nagy, 2002). Thus, we examined whether the trafficking of pPKC-like antigens also depends on the actin cytoskeleton. When interphase embryos were treated with Cytochalasin B, pPKC-like antigens did not move to the cortex, but remained close to the nucleus (Supplemental Fig. S6). In addition, the division axis was aligned randomly rather than with the apical cortex (Supplemental Fig. S6). These show that actin cytoskeleton is required for movement of pPKC-like antigens and proper orientation of division axis. These results also suggest that serotonin signaling could regulate changes in the actin cytoskeleton to polarize the cell. However, the actin cytoskeleton is not severely disrupted in metergoline-treated embryos, as these embryos completed cytokinesis in the 4-8 division and 8-12 division and displayed normal cortical phalloidin staining (Fig. 2). Therefore, it is more likely that serotonin signaling regulates events downstream of the organized cortical actin cytoskeleton.

### 2.5. A model for serotonin signaling in asymmetric cleavage

Our results are consistent with a model in which serotonin signaling in *Ilyanassa* regulates spindle orientation through G-protein regulation of both the actin and microtubule cytoskeleton. The serotonin receptors 1, 2 identified in early *Ilyanassa* embryos, are G-protein coupled receptors (GPCR). Experimental results in other model systems have shown and that GPCRs can activate a Gα subunit to recruit the PAR-complex to the apical cortex (Yoshiura et al., 2012) and that Gα and Gβ subunits can regulate spindle position and orientation (Gotta and Ahringer, 2001; Zwaal et al., 1996). Also, it has been shown that serotonin signaling regulates actomyosin contractility for cell intercalation during the *Drosophila* germband extension (Karki et al., 2023). In this connection, it is worthwhile to note that the actin cytoskeleton is required for proper spindle orientation in early mollusc embryos *Lymnaea* (Shibazaki et al., 2004) and *Ilyanassa* (this study, Supplemental Fig. S6). Based on these results, we propose a regulatory network in which a serotonin regulated GPCR recruits the microtubules of the spindle via polarity proteins and independently recruits a suite of scaffolding proteins that anchor the mitotic spindle via the actin cytoskeleton.

Interestingly, early *Ilyanassa* blastomeres separated from all other blastomere contacts at the two and four cell stages undergo normal division patterns, with essentially wildtype asymmetry and spindle orientation (Wandelt et al., 2023). Similarly, blastomere isolation experiments in other spirally cleaving embryos suggest that spindle orientation does not rely on sustained intercellular contacts (Costello, 1945; Wilson, 1904). Thus, the observed dependence of spindle polarity on serotonin signaling must require an intercellular signaling that occurs prior to blastomere isolation or is cell autonomous. An autocrine or intracellular function for serotonin signaling has not yet been documented. However, serotonin co-localizes with the mitotic apparatus in the early *Ilyanassa* embryo, and several lines of evidence suggest that functional serotonin receptors localize intracellularly in various cell types and embryos (Buznikov et al., 2001; Buznikov et al., 2003; Cornea-Hebert et al., 2002; Emanuelsson, 1992). Thus, it remains a formal possibility that each macromere has an intrinsic polarity at birth and intracellular serotonin signaling is required to maintain that polarity. Regardless of the mechanism, our experiments define a critical role for serotonin signaling in early asymmetric cleavage. Future experiments will be required to determine the extent to which this serotonin function is conserved in other spiralians, as well asymmetric cell divisions and stem cell division in other model systems.

## 3. Materials and Methods

### 3.1. Animals, immunohistochemistry, and Western blotting

Snail husbandry, immunohistochemistry, and Western blotting were done as previously described (Gharbiah, 2009; Lambert and Nagy, 2001). Anti-5-HT antibody (ImmunoStar), anti-tubulin antibody (E7, Developmental Studies Hybridoma Bank, University of Iowa) were used for immunohistochemistry. The anti-5-HT is known to react with serotonin in various species, including vertebrates (Hunt et al., 2005), arthropods (Pulver and Marder, 2002), and molluscs (Fickbohm et al., 2005). Anti-5-HT1 antibody (Gene Tex) was used for Western blotting. DAPI (Sigma) and phalloidin Alexa 488 (Sigma) were used to visualize the nucleus and actin cytoskeleton. In the pre-absorption control of the 5-HT antibody (ImmunoStar), the antibody was incubated with 5-HT-BSA conjugate (20 µg/ml, ImmunoStar) in phosphate buffer saline, 0.3% Triton, 2% BSA for 18-24 hours at 4°C.

### 3.2. Drug treatments

Metergoline (Sigma) was dissolved in EtOH at 90 mg/ml and diluted in filtered artificial seawater (FASW) to 20 µM or 30 µM. Arachidonyl serotonin (AA-5-HT, 20 mg/ml, Tocris) was diluted in FASW to 30 µM. Cytochalasin B (Calbiochem) was dissolved in DMSO at 20 mg/ml and diluted in FASW to 20 µg/ml. Controls received the same dilution of EtOH or DMSO.

### 3.3. RT-PCR analysis

RNA was isolated with Trizol (Invitrogen), and cDNA was synthesized with high-capacity RNA-to-cDNA Master Mix (Applied Biosystems) according to the manufacturer’s instructions. The nucleotide sequences of the primers are the following:

5-HT1-forward 5’-TGACCTCGCTCTTCCTGTG-3’,
5-HT1-reverse 5’-GCACACACAGCCAGATTATCA-3’,
5-HT2-forward 5’-GTAACGCTCCAGGCTGATG-3’,
5-HT2-reverse 5’-CTAGAGAAGCGGCTCCAGAA-3’,
40s rRNA-forward 5’-GAAGAAGCCGAAGTTTGACG-3’,
40s rRNA-reverse 5’-ATCACTGGTGGTTCGTAGCC-3’.

The amplification parameters were 94°C for 5 minutes, 40 cycles of 94°C for 30 seconds, 64°C for 30 seconds, 72°C for 1 minute, and a final extension at 72°C for 10 minutes.

### 3.4. Image analysis

ImageJ software was used for image analyses. Cell outlines were selected with the polygon selection tool to measure the size ratio of daughter cells using the phalloidin staining as a cortical marker. The area measurement tool was used to compare the size ratio of the selected cells. To measure the angle between pPKC-like antigens and the animal-vegetal axis, the center of the pPKC-like antigens was estimated by the centroid measurement tool, and the angle between the animal-vegetal axis and the center of pPKC-like antigens was determined with the angle tool.

### 3.5. Mass spectrometry

Proteins from the 4-cell embryos were isolated, as described previously (Gharbiah, 2009). The purified proteins were separated by SDS-PAGE, and the gel was stained with coomassie blue. The 40-45 kDa gel band was cut out and digested with trypsin. The digest mixture was separated by liquid chromatography-tandem mass spectrometry (LC-MS/MS). The spectra were searched against an NCBI non-redundant database or serotonin receptor sequences from various metazoan animals.

## Acknowledgements

The authors thank A. Wodarz and D. Zarnescu for Bazooka/PAR-3 antibody; B. Davidson, SD Hester, MM Stansbury, and lab members for critical manuscript reading; D. Lambert for sharing the 454 EST database. We thank Jessica Wandelt, James Cooley, Aldo Figueroa, and Dillon Steadman for their expert technical assistance and discussions. Mass spectrometric data were acquired by the Arizona Proteomics Consortium supported by NIEHS grant ES06694 to the SWEHSC, NIH/NCI grant CA023074 to the AZCC and by the BIO5 Institute of the University of Arizona. This study was supported by a grant from National Science Foundation (0820564) to L. M. N. This study was also partly supported by Grant-in-Aid for Scientific Research (KAKENHI) (22K06302 to A.N.), Japan Society for the Promotion of Science.

## Author contributions

Conceptualization: A.N., L.M.N.; Methodology: A.N., L.M.N.; Validation: A.N., L.M.N.; Formal analysis: A.N., L.M.N.; Investigation: A.N.; Resources: L.M.N.; Data curation: A.N., L.M.N.; Writing - original draft: A. N.; Writing - review & editing: A. N., L.M.N.; Visualization: A. N.; Supervision: A. N., L.M.N.; Project administration: A. N., L.M.N.; Funding acquisition: L.M.N., A.N.

---

**Supplemental Figure S1.**
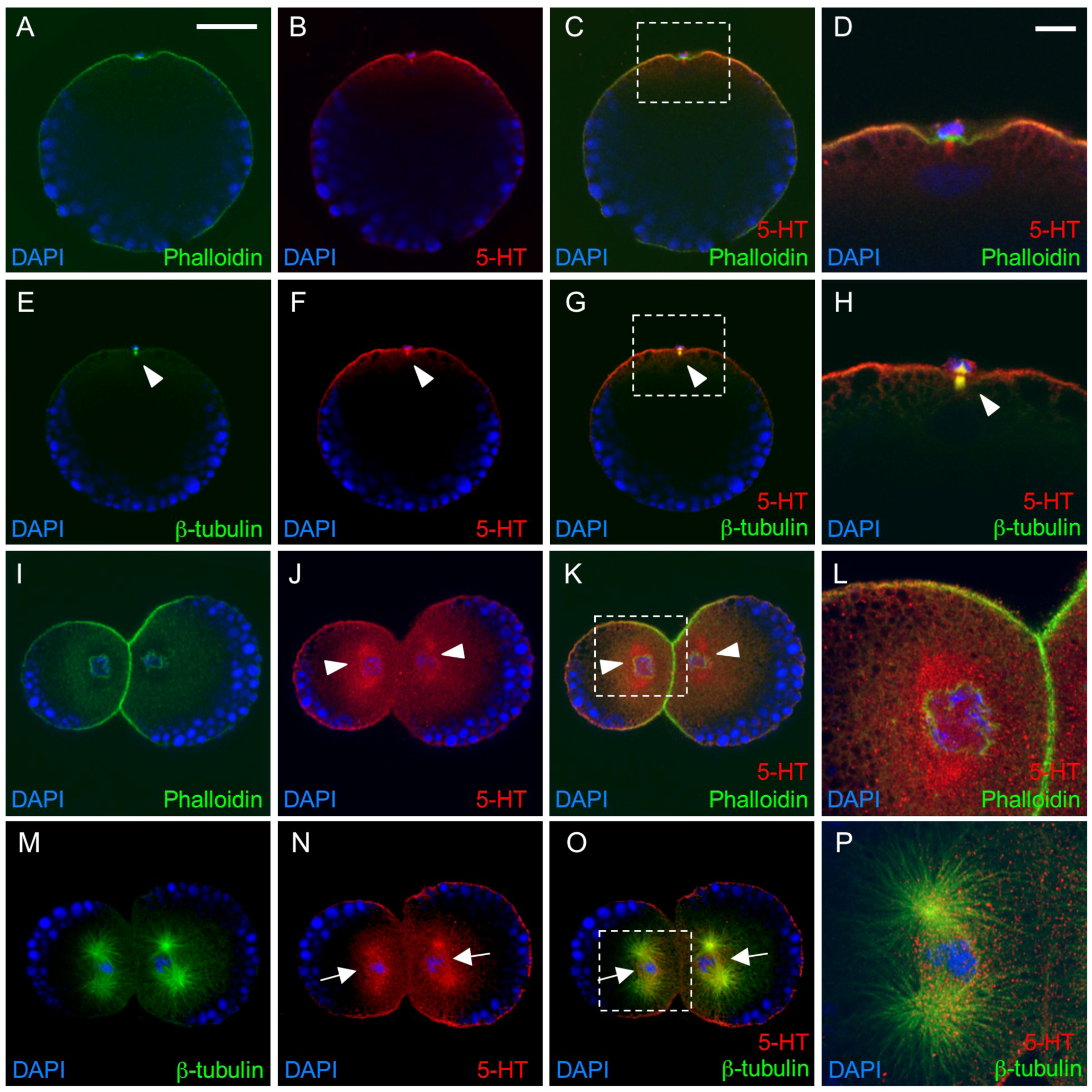
Expression of serotonin (5-HT) during 1- and 2-cell stages. (A-H) 1-cell stage, side view. The animal pole is toward the top. (I-P) 2-cell stage, apical view. The animal pole is toward the viewer. Nuclei were stained with DAPI (blue). Actin cytoskeleton was stained with phalloidin (green in A-D and I-L). β-tubulin (green in E-H and M-P) and serotonin (5-HT, red) were stained with specific antibodies. (D, H, L, P) Higher-magnification view of the inset of C, G, K, and O, respectively. In the 1-cell stage (A-H), serotonin was detected in the cell cortex of the animal hemisphere. Co-staining with β-tubulin antibody suggested that serotonin also localized to the meiotic spindle (E-H, arrowhead). In the 2-cell stage, serotonin is localized to the cell cortex (I-K). Serotonin was also concentrated around the interphase nuclei (J-L, arrowheads) and metaphase spindle (N-P, arrows). Scale bars, 50 µm (A-C, E-G, L-K, M-O); 10 µm (D, H, L, P).

**Supplemental Figure S2.**
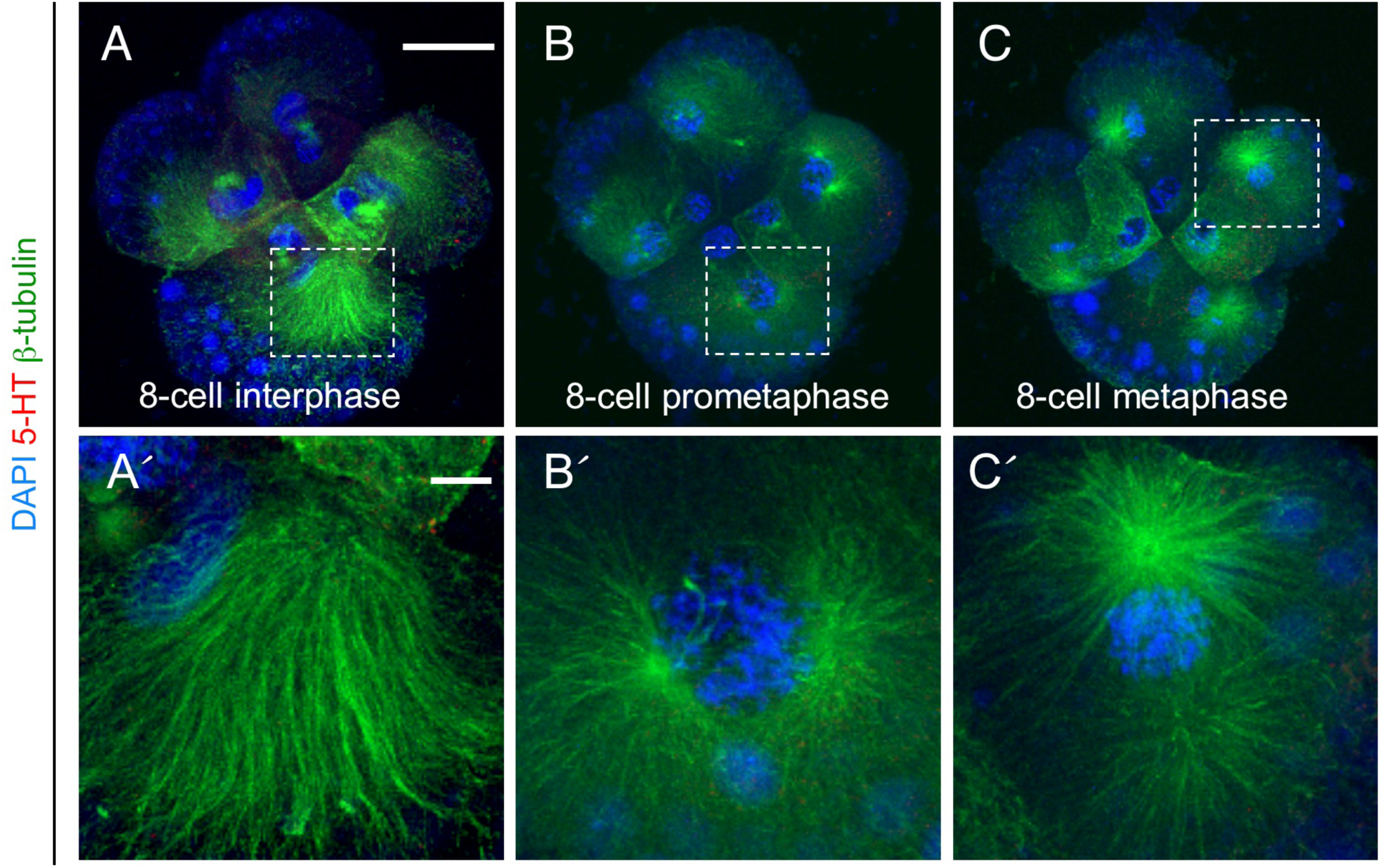
Preincubation of 5-HT antibody with 5-HT-BSA conjugate. No specific signal is detected in interphase (A), prometaphase (B), or metaphase (C) of the 8-cell stage. (A′-C′) Higher-magnification view of the inset of a, b, and c, respectively. Scale bars, 50 µm (A-C); 10 µm (A-C′).

**Supplemental Figure S3.**
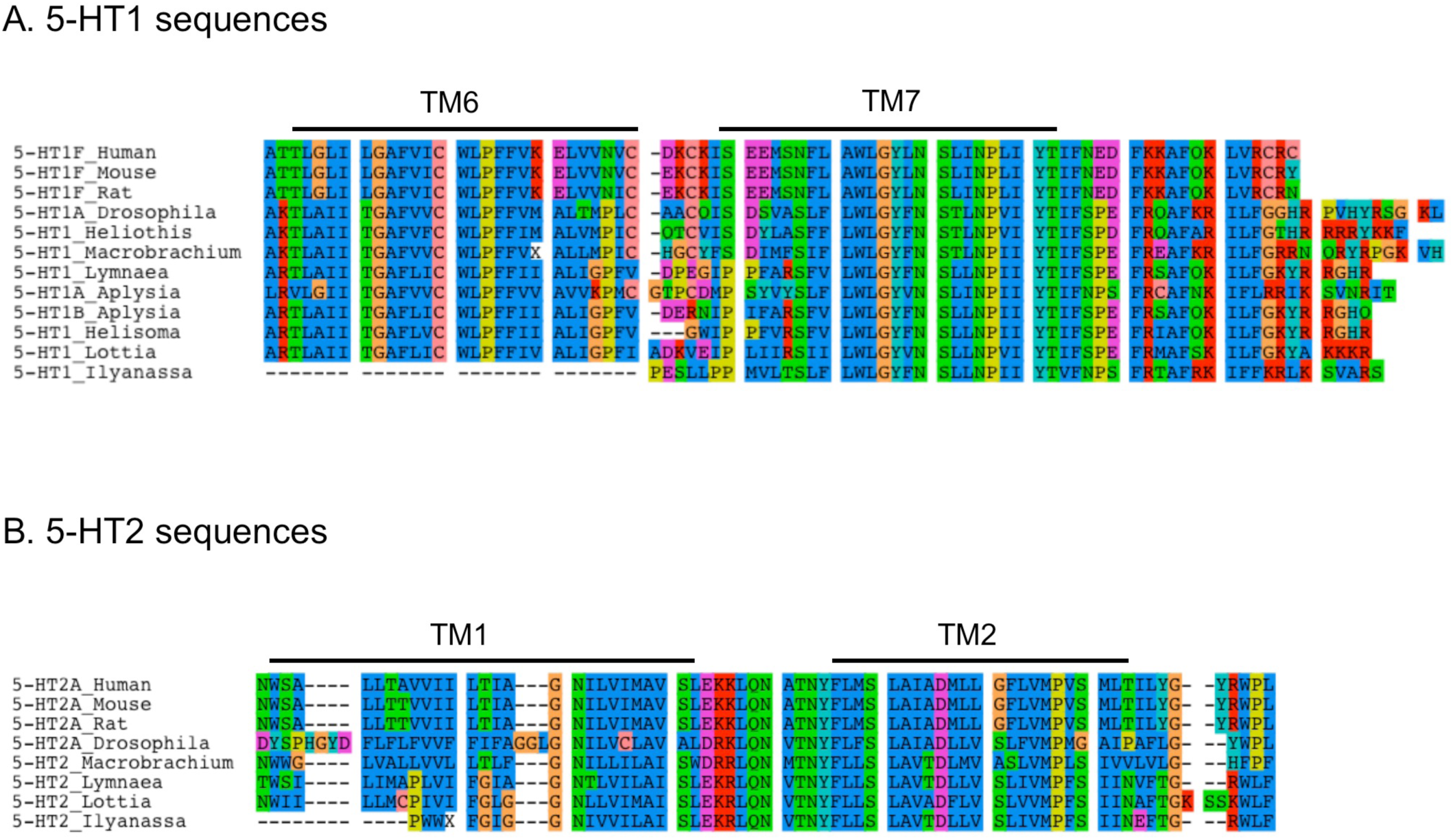
Alignment of amino acid sequences for 5-HT1 (A) and 5-HT2 (B) from various metazoans. Black lines indicate transmembrane regions (TM). Note that TM6 and TM7 of 5-HT1, which are recognized by the polyclonal antibody (Gene Tex), are highly conserved in metazoans. GenBank accession numbers are: 5-HT1F_Human (*Homo sapiens*, NP_000857.1), 5-HT1F_Mouse (*Mus musculus*, NP_032336.1), 5-HT1F_Rat (*Rattus norvegicus*, NP_068629.2), 5-HT1A_Drosophila (*Drosophila melanogaster*, NP_476802.1), 5-HT1_Heliothis (*Heliothis virescens*, CAA64863.1), 5-HT1_Macrobrachium (*Macrobrachium rosenbergii*, ACB38667.1), 5-HT1_Lymnaea (*Lymnaea stagnalis*, AAA29290.1), 5-HT1A_Aplysia (*Aplysia californica*, AAC28786.1), 5-HT1B_Aplysia (*Aplysia californica*, AAM46088.1), 5-HT1_Helisoma (*Helisoma trivolvis*, AAQ95277.1), 5-HT2A_Human (*Homo sapiens*, NP_000612.1), 5-HT2A_Mouse (*Mus musculus*, NP_766400.1), 5-HT2A_Rat (*Rattus norvegicus*, NP_058950.1), 5-HT2A_Drosophila (*Drosophila melanogaster*, NP_524223.2), 5-HT2_Macrobrachium (*Macrobrachium rosenbergii*, ABM01873.1), 5-HT2_Lymnaea (*Lymnaea stagnalis*, AAC16969.1). In *Lottia gigantea,* orthologs of 5-HT1 and 5-HT2 were identified in the genome sequences in the JGI (DOE Joint Genome Institute; http://genome.jgi-psf.org/Lotgi1/Lotgi1.home.html). The protein IDs of Lottia_5-HT1 and Lottia_5-HT2 are 134511 and 82265, respectively.

**Supplemental Figure S4.**
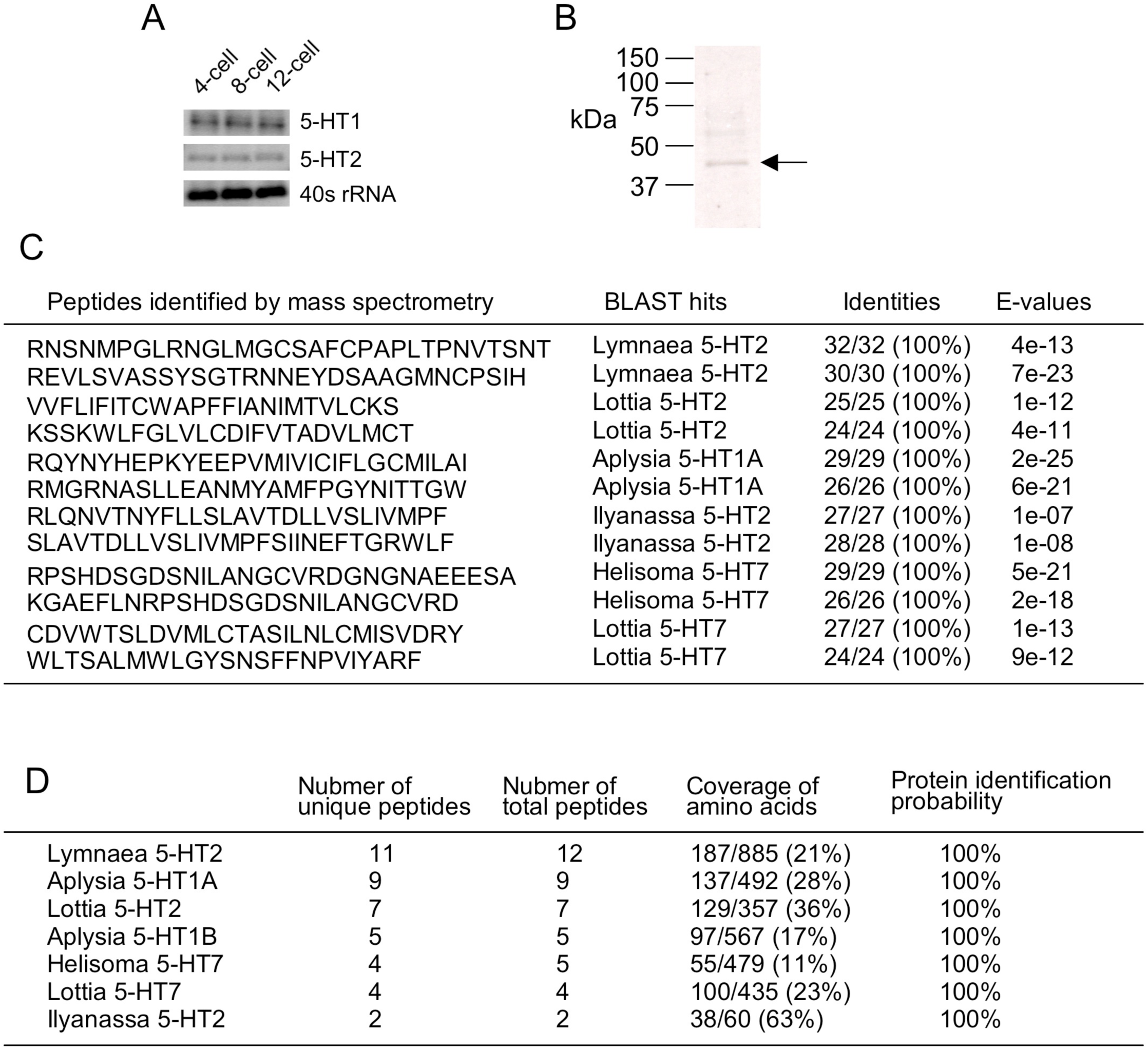
Expression of serotonin receptors in early *Ilyanassa* embryos. (A) RT-PCR shows the expression of the transcripts of serotonin receptors 5-HT1 and 5-HT2 during the early cleavage stages. 40s RNA was used as a positive control. (B) Western blotting with anti-serotonin receptor antibody. A 40-45 kDa band, similar in size to other serotonin receptors of vertebrates, is detected in the 4-cell embryo. (C) Examples of identified peptides from the 40-45 kDa gel band. Peptides that belong to molluscan serotonin receptors are shown. (D) Summary of the number of identified peptides and their amino acid coverage. For example, 11 unique peptides (a total of 12 peptides) have significant similarities with *Lymnaea* 5-HT2, and their amino acid coverage is 187/885 (21%). Protein identification probabilities were calculated using the software Scaffold version 3. Note that partial sequences (60 amino acids) are identified in *Ilyanassa* 5-HT2 (Supplemental Figure S3).

**Supplemental Figure S5.**
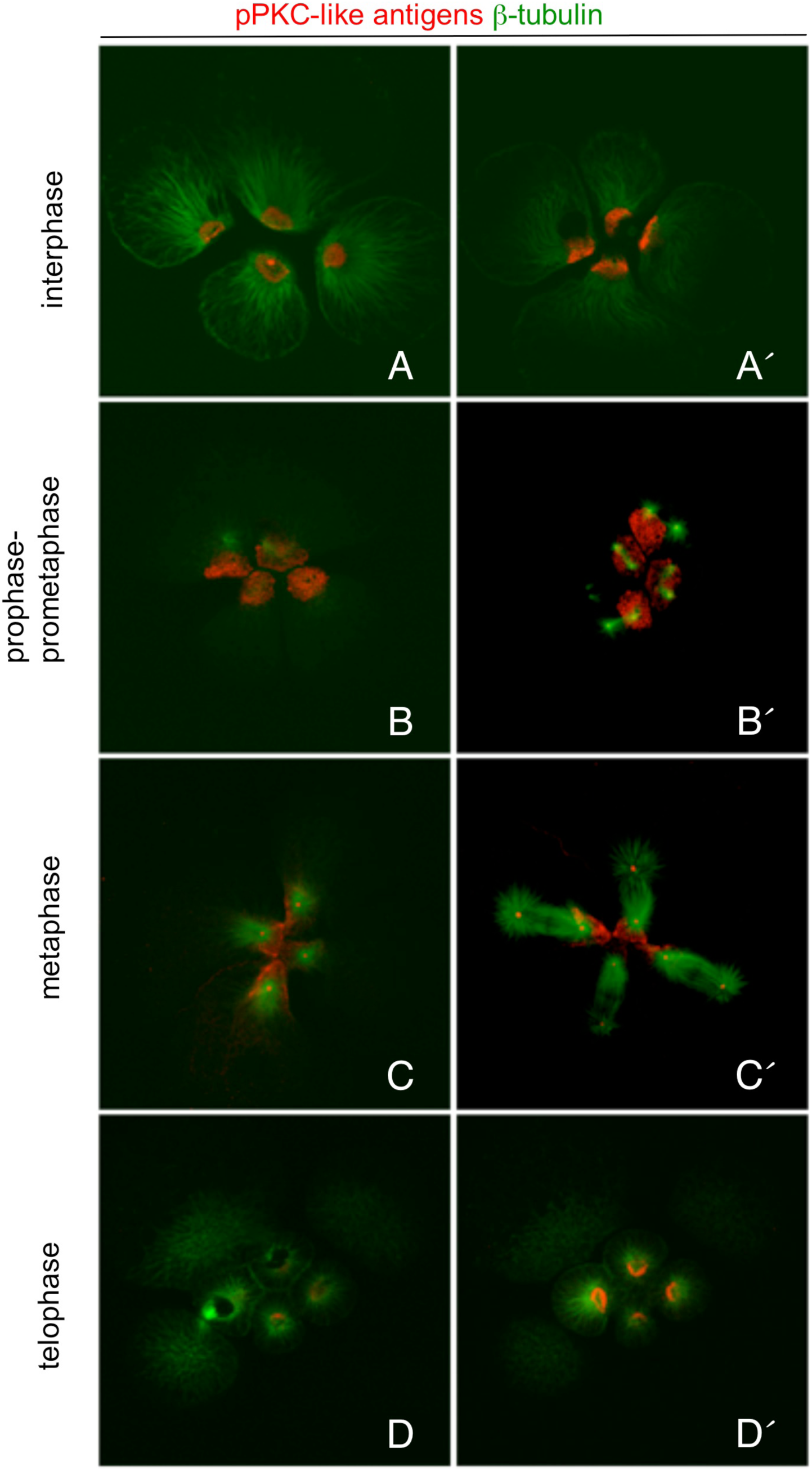
**Localization** of pPKC-like antigens is cell cycle-dependent. Embryos were fixed throughout the 4- to 8-cell stage and assayed for the expression of pPKC-like antigen. pPKC-like antigen (red) and beta-tubulin (green) were stained with their specific antibodies. (A, A′) Different confocal sections of an interphase-prophase embryo where pPKC-like antigens are expressed in the center of the interphase microtubule array. (B, B′) Different confocal sections of a prophase-prometaphase embryo showing pPKC-like antigens localized to the anterior cortex of each macromere. The mitotic spindle is forming, and pPKC-like antigens are also localized in the center of the microtubule array at each end of the spindle. (C, C′) Different confocal sections of a metaphase embryo. pPKC-like antigens are localized at the anterior cortex of each macromere and are also present at the center of both ends of all four mitotic spindles. The anterior end of the mitotic spindle is focused on the patch of pPKC-like antigens at the anterior cortex (C). (D, D′) Different confocal sections of a telophase embryo show the halos of pPKC-like antigens in the four daughter micromeres.

**Supplemental Figure S6.**
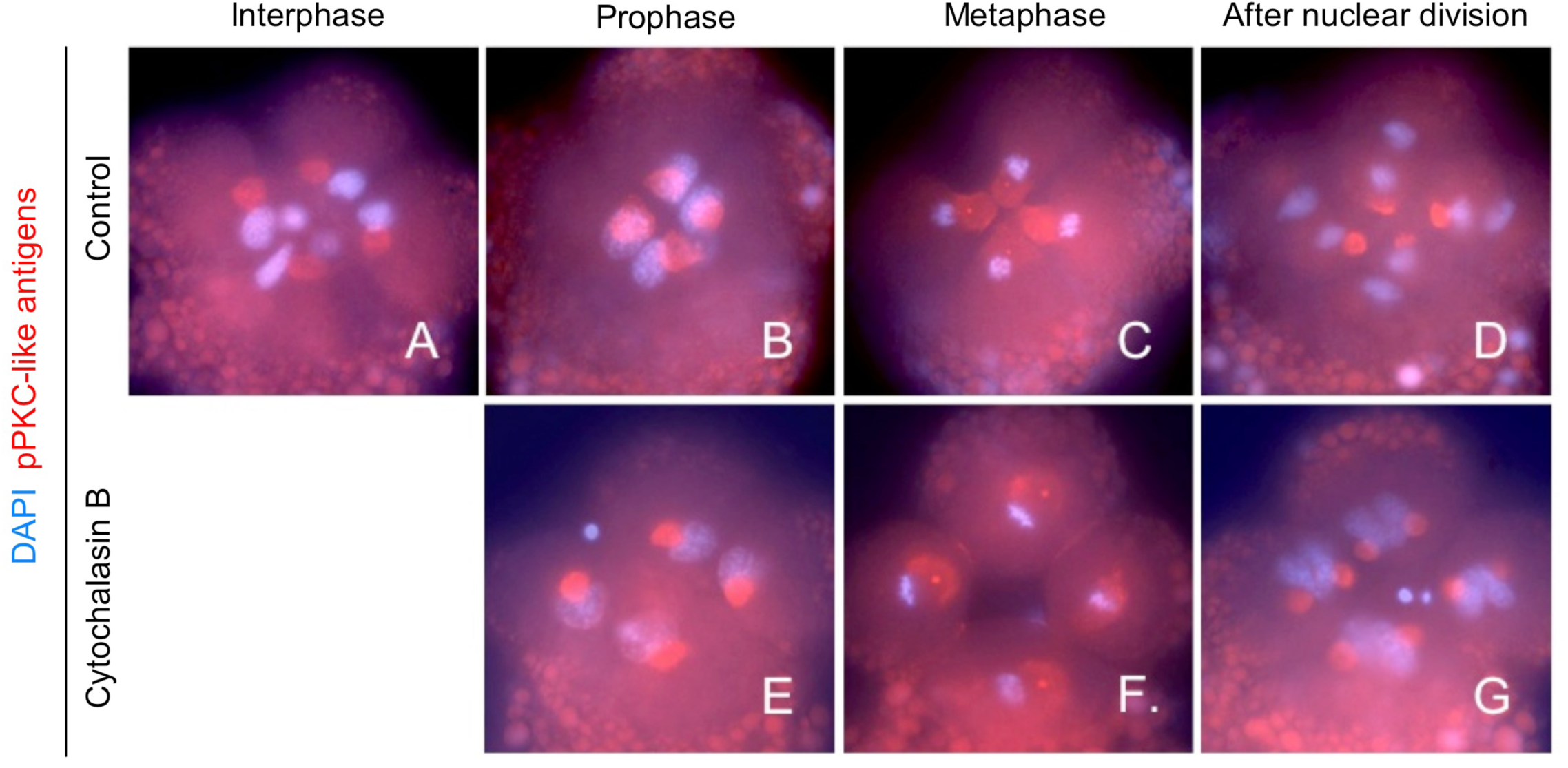
Microfilaments are required to move pPKC-like antigens from the centrosome to the cortex. To determine the role of microfilaments in the movement of pPKC-like antigens from the centrosome to the cortex, embryos were incubated in Cytochalasin B (CB) starting at interphase when pPKC-like antigens are localized on the centrosome (A). (A-D) Expression of pPKC-like antigens (red) in control embryos. (A) pPKC-like antigens localize to the centrosome in interphase cells. (B) At prophase, pPKC-like antigens are localized to the anterior cortex of each macromere. (C) pPKC-like antigens are localized to the anterior cortex and each spindle pole at metaphase. (D) After division, pPKC-like antigens are localized on the interphase centrosome of all eight blastomeres. (E-G) Expression of pPKC-like antigens in CB-treated embryos. (E) At prophase in CB-treated embryos, pPKC-like antigens are still localized to a spherical structure. pPKC-like antigens are localized in a non-cortical patch in each macromere by metaphase. pPKC-like antigens are also detected on each spindle pole. The orientation of each spindle is random. (G) When the nuclei have completed division, pPKC-like antigens are localized on the centrosome adjacent to each nucleus. Note the irregular arrangement of the nuclei. Red: pPKC-like antigens; blue: DAPI.

